# Inferring the role of habitat dynamics in driving diversification: evidence for a species pump in Lake Tanganyika cichlids

**DOI:** 10.1101/085431

**Authors:** Thijs Janzen, Rampal S. Etienne

## Abstract

Geographic isolation that drives speciation is often assumed to slowly increase over time, for instance through the formation of rivers, the formation of mountains or the movement of tectonic plates. Cyclic changes in connectivity between areas may occur with the advancement and retraction of glaciers, with water level fluctuations in seas between islands or in lakes that have an uneven bathymetry. These habitat dynamics may act as a driver of allopatric speciation and propel local diversity. Here we present a parsimonious model of the interaction between cyclical (but not necessarily periodic) changes in the environment and speciation, and provide an ABC-SMC method to infer the rates of allopatric and sympatric speciation from a phylogenetic tree. We apply our approach to the posterior sample of an updated phylogeny of the *Lamprologini*, a tribe of cichlid fish from Lake Tanganyika where such cyclic changes in water level have occurred. We find that water level changes play a crucial role in driving diversity in Lake Tanganyika. We note that if we apply our analysis to the Most Credible Consensus (MCC) tree, we do not find evidence for water level changes influencing diversity in the *Lamprologini*, suggesting that the MCC tree is a misleading representation of the true species tree. Furthermore, we note that the signature of habitat dynamics is found in the posterior sample despite the fact that this sample was constructed using a species tree prior that ignores habitat dynamics. However, in other cases this species tree prior might erase this signature. Hence we argue that in order to improve inference of the effect of habitat dynamics on biodiversity, phylogenetic reconstruction methods should include tree priors that explicitly take into account such dynamics.

## INTRODUCTION

Environmental changes such as the formation of mountain ridges, the formation of rivers and the movement of tectonic plates have long been known to be important drivers of speciation (Coyne and Orr 2004). Repeated environmental changes may thus lead to diversification patterns. Cyclic changes in the environment can cause populations to continuously switch between an allopatric and sympatric stage, providing a continuously renewed potential for speciation. And these cyclic changes can in turn drive diversity towards levels unexpected given the current geography, sometimes referred to as a “species pump” (Heaney 1985; Rossiter 1995). Examples of species pumps include environmental fluctuations fragmenting habitats on the slopes of mountains (Weir 2006; Sedano and Burns 2010; Hutter et al. 2013), glaciations and postglacial secondary contacts (Barnosky 2005), sea level changes causing the fusion and fragmentation of islands (Glor et al. 2004; Thorpe et al. 2008, but see Papadopoulou and Knowles 2015), and fluctuations in water level causing fragmentation and fusion of lakes with uneven bathymetry, as in the African Rift Lakes (Cohen et al. 1997b; Alin et al. 1999; McGlue et al. 2008; Ivory et al. 2016).

The African Rift Lakes provide a good starting point in studying the interplay between cyclic habitat dynamics and speciation, because they have been subject to frequent water level changes (Cohen et al. 1997b; Alin and Cohen 2003; Ivory et al. 2016), and are well known for their tremendous biodiversity (Seehausen 2000, 2006; Turner et al. 2001; Wagner et al. 2012, 2014; Brawand et al. 2014). An estimated number of 2000 cichlid fish species (Turner et al. 2001) have evolved in the African Rift Lakes over the past 10 million years (Genner et al. 2007; Meyer et al. 2016), and comprise one of the most spectacular adaptive radiations (Seehausen 2006). The most prominent water level changes took place in Lake Tanganyika, where the water level has dropped substantially on multiple occasions over the past million years, sometimes splitting the lake into multiple smaller lakes (Lezzar et al. 1996; Cohen et al. 1997a, 2007). Being the oldest lake of the three large rift lakes (Cohen et al. 1993), Lake Tanganyika contains the highest behavioral diversity (Konings 2007) and is the only lake with a highly resolved phylogeny for cichlid fish. Evidence for the influence of changing water levels comes from analysis of mitochondrial DNA, which shows that for *Tropheus* species, some populations have experienced secondary contact upon changes in water level, potentially increasing genetic diversity and driving speciation (Sturmbauer et al. 2001; Koblmüller et al. 2011; Sefc et al. 2017). Similar patterns were found for *Variabilichromis moorii* and *Ophthalmotilapia nasuta* (Sturmbauer et al. 2001), *Telmatochromis temporalis* (Winkelmann et al. 2016), and *Altolamprologus* (Koblmüller et al. 2016). Comparison of mitochondrial DNA between populations from deep and shallow areas emphasizes that the deep areas are habitats that are more persistent over time, with lower genetic variation (Nevado et al. 2013). Furthermore, *Eretmodus* lineages identified using mitochondrial DNA are strongly associated with the bathymetric basins of Lake Tanganyika (Verheyen et al. 1996), suggesting that they have independently diversified at low water level.

Aguilée et al. (2013) developed a model for the African Rift Lakes in which populations at different locations diverge from each other depending on the local habitat, and at the same time allowed for sympatric speciation by implementing assortative mating that allows for a single branching point in trait values. Over time the different locations become separated or are reconnected, and this may drive the formation of new species. The authors conclude that stable numbers of diversity are best obtained by a fragmented habitat with recurrent merged states and rapid fluctuations. However, Aguilée et al. (2013) do not compare their results to empirical data. By contrast, Pigot et al (2010) used a spatially explicit model of landscape fragmentation, where consecutive splitting of species’ geographic ranges drives speciation, and compared phylogenies generated with their model, with known bird phylogenies. They found that including this geographical context of speciation explains a large part of the features exhibited by the reconstructed avian trees. Hence, including a geographical context of speciation seems a promising research avenue.

Here, we provide a method to infer whether or how cyclic changes in the environment influence both the generation and the maintenance of biodiversity. We use an extension of the standard constant-rates birth-death model. Because deriving an expression for the likelihood of this model for a given set of phylogenetic branching times is difficult, but simulation of phylogenies under the model is easy, we used approximate Bayesian computation (ABC) based on sequential Monte Carlo sampling (SMC) to estimate parameters from phylogenies. We applied our approach to an updated phylogeny of the *Lamprologini*, a tribe of cichlid fish from Lake Tanganyika in order to assess the importance of these habitat dynamics in shaping the current biodiversity of cichlids in Lake Tanganyika.

## METHODS

### Model

To model the interaction between environmental change and speciation, we envisage a lake that consists of a single pocket at high water level, but that splits into two pockets when the water level drops. When the water level drops, we assume that all species distribute themselves equally over the two pockets; similarly, when the water level rises, all species previously contained in the two pockets are combined into the single pocket. Allopatric speciation can only occur when the water level is low. We assume a constant probability rate for allopatric speciation, and hence the waiting time until the next speciation event is exponentially distributed. After this waiting time, one of the two incipient species in either pocket can speciate into a new species. If this allopatric speciation does not occur before the water level rises again, i.e. reflecting that there has not been enough genetic divergence, the two incipient species in the two pockets merge back into one species. This is conceptually similar to the idea of protracted speciation (Etienne and Rosindell 2012): the water level drop initiates the speciation process whereas the allopatric speciation event is the completion of speciation under the protracted speciation model. Sympatric speciation can always occur in our model, either at high water level in the lake, or in both pockets when the water level is low. Extinction is considered to be a background process that occurs locally, i.e. within a pocket. If the water level is high, this causes extinction of a species, if the water level is low, this causes local extinction in one of the pockets.

We implemented our model using a Gillespie algorithm, where the time steps are chosen depending on the rate of possible events. In the model there are five possible events:

1. A water level change event, inducing incipient species or merger of incipient species.
2. Sympatric speciation event at high water level, with rate 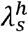
3. Sympatric speciation event at low water level, with rate 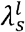
4. Allopatric speciation(-completion) event, at low water level, with rate 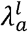
5. Extinction event, with rate 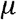

When the water level drops, all species distribute themselves over both pockets. Thus, immediately after a water level drop, the number of incipient species is equal to twice the number of species. When the water level rises, all incipient species that belong to the same species merge into a single species. During a sympatric speciation event, a single species splits into two new species, and the original (incipient) species is consumed in the process. Here we assume that local disruptive selection causes divergence, similar to the implementation of speciation by Aguilee (Aguilée et al. 2011, 2013). If sympatric speciation occurs when the water level is low, the species in the other pocket is retained, and thus three new lineages arise: the first branching point occurs at the water level drop while the second occurs at the sympatric speciation event (Figure 1).

**Figure 1.**
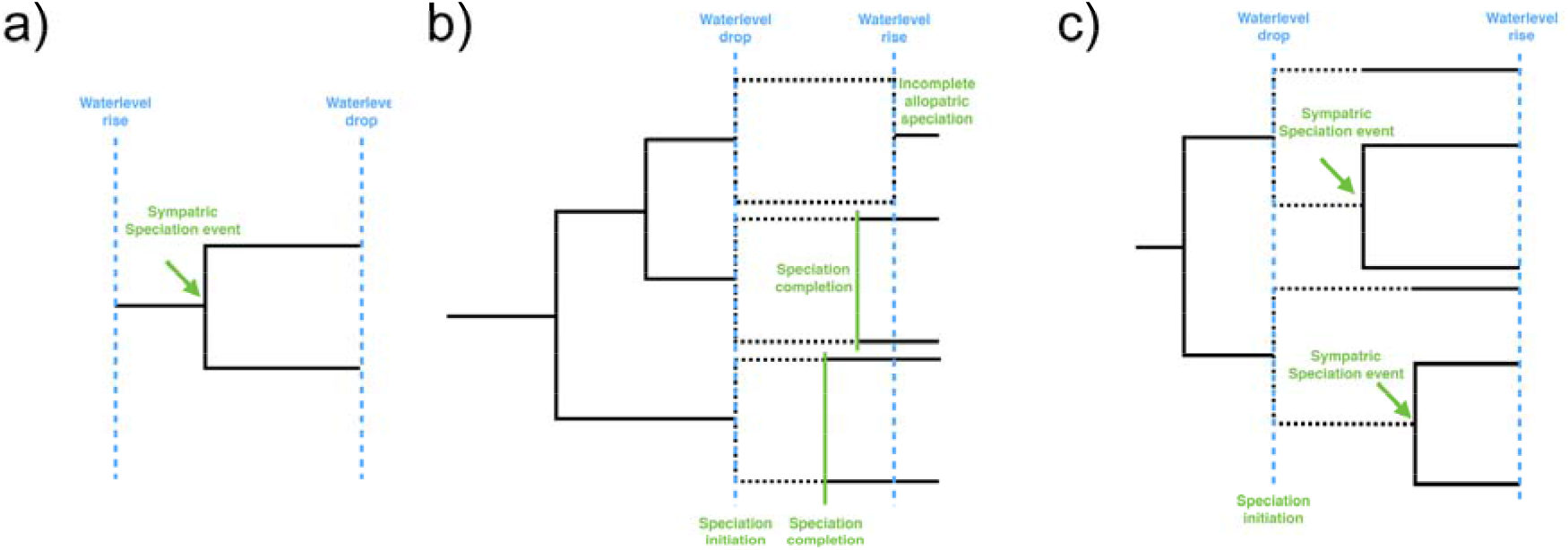
Schematic representation of the consequences of the three different types of speciation. Time proceeds from left to right. The dotted blue lines indicate water level changes. (a) During a sympatric speciation at high water level event, diversification is not aligned with any associated water level change. (b) During allopatric speciation at low water level, speciation initiation (incipient species are indicated with a dotted line) coincides with the water level drop, causing branching events (if speciation-completion occurs before water level rise) to line up in time. Branching events are conditionally independent of the time of speciation completion, hence, even when the actual speciation completion events occur at different time points, branching events in the species tree are identical. (c) During a sympatric speciation event at low water, the speciation event is independent of the water level changes. Because the original species is consumed in the process, a new branching event is also added at the water level change event. Hence, both speciation-completion (b) and sympatric speciation at low water level (c) cause branching times to line up at the time of water level drop. Please note that (a) and (c) represent the reconstructed species tree, but (b) does not; the reconstructed species tree for (b) would not show the branching event in the uppermost part of the tree.

**Figure 2.**
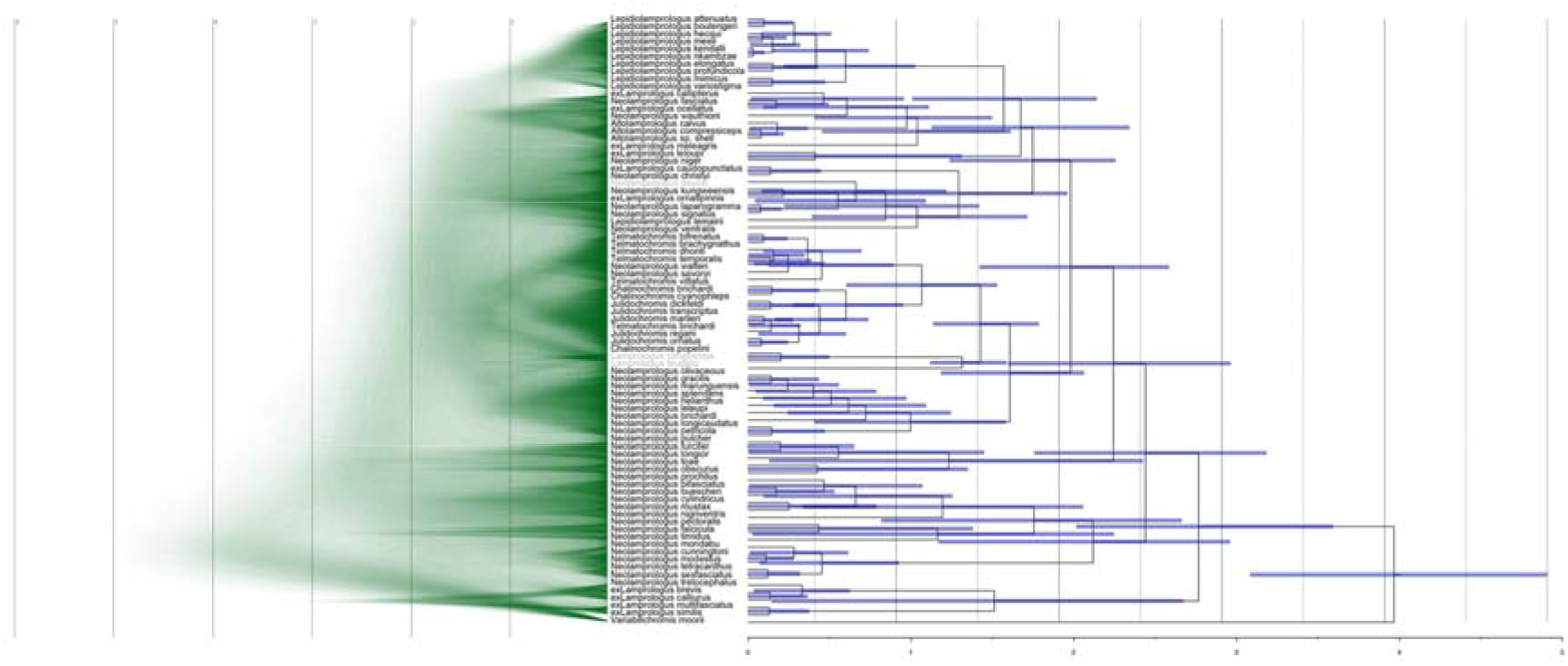
Phylogenetic hypothesis for the *Lamprologini* and outgroups, based on 3 mitochondrial and 6 nuclear genes, and two calibrations: 4 million years for the root of the *Lamprologini* clade, and 1.1 - 3.5 million years for the Congo *Lamprologini* species. Left panel: DensiTree (Bouckaert and Heled 2014) representation of the MCMC chain obtained using *BEAST. Shown are trees from a thinned posterior chain, after selecting every 100,000^th^ tree. Riverine species are indicated in grey. Right panel: Maximum Clade Credibility tree. Bars around the node span the 95% HPD for each node. Please note that for the dual display of both the densitree representation and the MCC phylogeny, some tips of the MCC phylogeny might appear slightly misaligned. A high resolution version of both the Densitree representation and the MCC phylogeny can be found in the supplementary information.

During an extinction event, one (possibly incipient) species is removed from the simulation. If the water level is low, this need not lead to the extinction of a species, because the sister incipient species might remain in the other pocket, ensuring survival of the species.

### Maximum Likelihood

Without water level changes, our model reduces to the constant rates birth-death model (Nee et al. 1994). As a reference therefore, we estimated parameters of the standard birth-death model using Maximum Likelihood. The likelihood of the birth-death model was calculated using the function “bd_ML” from the R package DDD. (Etienne et al. 2012).

### Fitting the model to empirical data

We performed two different fitting procedures: firstly, we performed a model selection procedure, where three different water level scenarios were fitted simultaneously to the data (more information about the chosen scenarios can be found in the next section). The model selection procedure simultaneously estimates parameter estimates and assesses the fit of the models. However, because the model selection procedure primarily samples the best fitting model (by design), it does not allow for the comparison of parameter estimates across different models. Therefore, we also fitted the three different water level scenarios independently to the empirical data, and obtained posterior distributions for the parameters relevant to these scenarios.

We fitted our model to 100 trees randomly sampled from the MCMC chain obtained from the *BEAST analysis (see below), and to the Most Credible Consensus (MCC) tree.

#### Water level scenarios

The main focus of our approach is to assess the impact of water level changes on the diversification rate. Lake Tanganyika experienced low water level stands 35 - 40 k years ago (kya) (-160 meter), 169 - 193 kya (-250 meter), 262 - 295 kya (-350 meter), 363 - 393 kya (-350 meter) and 550 - 1100 kya (-650 – 700 meter) (Lezzar et al. 1996; Cohen et al. 1997a). The southern and northern basin of Lake Tanganyika are separated from each other by a ridge at a depth of 500 meter below present level. Although some of these water level changes may not have split up the lake completely, we assume here that these water level changes still caused sufficient disruption of migration between the northern and southern basin, to be equivalent to physical separation. Consequently, high water levels occurred between 0 – 35 kya, 40 – 169 kya, 193 – 262 kya, 295 – 363 kya and 393 - 550 kya. Unfortunately the geological record does not reveal whether any low water level stands occurred beyond 1.1 million years ago (Ma). This leaves us with two alternative scenarios: either no low water level stands occurred beyond 1.1 Ma, or these low water level stands have not been preserved accurately in the geological record.

In order to capture these two scenarios we performed inference using two alternative water level implementations. Firstly we used the exact literature values, assuming a high water level stand until 1.1 Ma. We refer to this scenario as LW (Literature Waterlevels). Secondly we assumed that before 1.1. Ma, water level changes occurred at the same *average* rate of water level change in the most recent 1.1 million years. In the recent 1.1 million years, the lake experienced 5 high water level stands, and 5 low water level stands, which amounts to 10 water level changes in total. To extrapolate water level changes to more than 1.1 Ma, we drew waiting times until the next water level change from an exponential distribution with rate 10. We refer to this scenario as EW (Extrapolated Water levels). Thirdly we also tested the null expectation: no effect of water level changes on speciation, we refer to this scenario as NW (No Water levels). Without water level changes, the model reduces to the constant-rates birth death model.

#### Parameter estimation

To fit the model to empirical data we used Approximate Bayesian Computation, in combination with a Sequential Monte Carlo scheme (ABC-SMC) (Toni et al. 2009). As summary statistics for the ABC analysis we chose the normalized Lineages Through Time statistic (Janzen et al. 2015), tree size, Phylogenetic Diversity (AvPD, Schweiger et al. 2008) and the *γ* statistic (Pybus and Harvey 2000). On all parameters 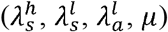 we chose uniform priors U(-3, 2), on a ^10^log scale, such that the eventual prior distribution spans (10^−3^, 10^2^). A ^10^log scale was chosen to explore parameter space uniformly, and put extra emphasis on low values. The standard deviation of the normal distribution used to perturb the parameters was chosen to have a mean of 0, and a standard deviation of 0.05 (on the ^10^log transformed parameter), and we updated one parameter each time (e.g. jumps were only made in one dimension, to avoid extremely low acceptance rates). The number of particles used per SMC step was 10,000, where a particle is a data structure containing the model choice and the parameter estimates. To assess the fit of the model to the data we calculated the Euclidian distance between the summary statistic of the simulated data and the empirical data. To ensure that the differences in summary statistics were on the same scale, we normalized the differences. Differences were normalized by dividing each difference by the standard deviation of that summary statistic of 1,000,000 trees simulated using parameter values sampled from the prior.

#### Model selection

To identify which model best explains the data, we performed ABC model selection, as described in Toni et al. (2009; 2010). The main difference between standard ABC-SMC and ABC-SMC including model selection is that the latter adds one parameter, which keeps track of the model. As jumping kernel between models we assumed a 50% probability of staying at the same model, and a 25% probability of jumping to either other model. We assumed a uniform prior across all three models; this translates to a probability of 1/3 for each model in the first iteration of the ABC-SMC procedure, and hence an expected number of 3333 particles assigned to each model in the first iteration. This reversible jump ABC-SMC model selection procedure results in a posterior distribution over the three models, where the model with most support is the model selected most across all particles. We can calculate the Bayes factor by taking the ratio of the number of particles assigned to the respective models (Toni et al. 2009). For example, the Bayes factor of LW/EW is the number of particles assigned to the model with literature water level changes divided by the number of particles assigned to the model with extrapolated water level changes. Because a model can receive zero particles, we set the Bayes factor for each model compared to the model with zero particles to the maximum support possible, which is the total number of particles: 10,000. To calculate the posterior support for a model, we calculate 2 ln (Bayes factor), following Kass and Raftery (1995). A transformed Bayes factor over a value of 2 then corresponds to substantial support for the considered model (Kass and Raftery 1995).

#### Model selection validation

To assess whether our ABC-SMC method can accurately infer the correct model, we simulated 100 datasets for each model (NW, LW & EW), with parameter values drawn from the prior. We report the median Bayes factor across the 100 replicates. If our method can accurately infer the correct model, we expect the median Bayes Factor (after 2 ln transformation) to be above 2 when comparing posterior support for the model with which the data was simulated to the other two models.

#### Measurement uncertainty

A phylogeny generated with a high rate of allopatric speciation and a high rate of water level changes tends to have multiple speciation events that are aligned in time (Figure 1, b). This is due to the fact that the onset of speciation is given by the time of water level change. Phylogenetic reconstruction methods such as BEAST (Bouckaert et al. 2014) currently do not allow for simultaneous branching events. Hence, when fitting the model, trees are generated that are by definition dissimilar from the empirical tree constructed using BEAST, even if underlying events are close to the original events. To circumvent this we perturbed the branching time of each node in the trees simulated using our model. In this way speciation events that were previously aligned in time now occur on slightly different time points, as in a tree from a BEAST analysis. We perturbed branching times by adding a random number drawn from a truncated normal distribution with mean 0, standard deviation *σ*, truncated by the minimum distance to either the daughter or the parent species. If there were no daughter lineages present, and the node gave rise to an extant species, the normal distribution was truncated to the minimum distance to the parent or the present time. Nodes were perturbed from past to present (leaving the crown in place, to ensure a phylogenetic tree with the same age as the empirical tree). The standard deviation of the perturbation kernel was included as an extra parameter to be inferred, with a uniform prior on (10^−3^,10^0^).

#### Empirical data

We fitted our model to the phylogenetic tree of the tribe of *Lamprologini*, the most diverse tribe within Lake Tanganyika, containing 79 species of cichlids in Lake Tanganyika (Day et al. 2007; Koblmüller et al. 2007; Sturmbauer et al. 2010). The *Lamprologini* are endemic to Lake Tanganyika and its surrounding rivers and all species are substrate brooders with shared paternal and maternal care. In contrast to the mouthbrooding species from the *Haplochromini*, the *Lamprologini* show little sexual dimorphism and dichromatism, which are well-known indicators for sexual selection (Kraaijeveld et al. 2011). We therefore expect that the *Lamprologini* is a good candidate for picking up signals from water level changes.

We reconstructed a new *Lamprologini* tree following the workflow of the most complete *Lamprologini* tree to date, which is a consensus tree based on the mitochondrial ND2 gene (Sturmbauer et al. 2010), but we added three newly described species *(Lepidiolamprologus mimicus* (Schelly et al. 2007), *Neolamprologus timidus* (Kullander et al. 2014b) and *Chalinochromis cyanophleps* (Kullander et al. 2014a)). Using phyloGenerator (Pearse and Purvis 2013), we downloaded sequences from GenBank for nine genes (GenBank access numbers can be found in the Supplementary Information). Genes were selected on the basis of species coverage (at least 25% of the 79 Lamprologini species for which molecular data is available), and whether or not the gene was crucial for inclusion of a species (e.g. for a number of species, the only available gene was ND2). After selection, our full dataset consisted of three mitochondrial genes: the NADH dehydrogenase subunit 2 (ND2 gene, sequences from Kocher et al. 1995; Klett and Meyer 2002; Clabaut et al. 2005; Duftner et al. 2005; Schelly et al. 2006; Day et al. 2007; Koblmüller et al. 2007, 2016; Schwarzer et al. 2009; Wagner et al. 2009; Sturmbauer et al. 2010; O’Quin et al. 2010; Kullander et al. 2014b; Weiss et al. 2015). The cytochrome b (cytb) gene (sequences from Salzburger et al. 2002; Nevado et al. 2009; Wagner et al. 2009; O’Quin et al. 2010; Matschiner et al. 2011, 2016; Kullander et al. 2014b; Shirai et al. 2014) and the cytochrome c oxidase subunit I (COI gene, sequences from Sparks and Smith 2004; Nevado et al. 2013; Kullander et al. 2014a, 2014b; Breman et al. 2016; Matschiner et al. 2016) and six nuclear genes: the nuclear locus 38A (38A, sequences from Muschick et al. 2012; Meyer et al. 2016), the 18S ribosomal RNA internal-transcribed spacer 1–2 with 5.8S and 28S ribosomal RNA partial sequences (18S, sequences from Nevado et al. 2009; Koblmüller et al. 2016), the recombinase activating protein 1 (rag1, sequences from Clabaut et al. 2005; Nevado et al. 2009; Kullander et al. 2014b; Shirai et al. 2014; Koblmüller et al. 2016; Meyer et al. 2016), the endothelin receptor B1 gene (ednrb1, sequences from Muschick et al. 2012; Santos et al. 2014), the ribosomal protein S7 (rps7, sequences from Schelly et al. 2006; Meyer et al. 2016)) gene and the rod opsin gene (RH1, sequences from Sugawara et al. 2002; Spady et al. 2005; Nagai et al. 2011; Meyer et al. 2015). GenBank access numbers for the used sequences can be found in the supplementary material.

Sequences were aligned using MAFFT (setting: --auto) (Katoh and Standley 2013), and subsequently, sequences were cleaned using trimAI (sites with >80% data missing were removed, e.g. setting −gt 0.2) (Capella-Gutiérrez et al. 2009). Rather than concatenating the alignments, we partitioned the data into subsets with independent sequence evolution models, which is more suitable for a dataset which is expected to show incomplete lineage sorting or hybridization (Meyer et al. 2016). To prepare alignments for use with partitionFinder, alignments were combined using SequenceMatrix 1.8 (Vaidya et al. 2011). The best partitioning found by partitionFinder 2.1.1 (Lanfear et al. 2012, 2016), partitioned the data into 5 subsets (unlinked branches, AICc selection criterion), with all nuclear genes into one subset (Rps7, ednrb1, 38A, 18S, RAG1 and RH1), with substitution model HKY+I+Γ. The remaining three mitochondrial genes (ND2, COI and cytb) were placed in separate subsets, each with a GTR+I+ Γ substitution model.

Using *BEAST (Heled and Drummond 2010) within the BEAST 2 package (Bouckaert et al. 2014), we inferred the time-calibrated species tree. We used an uncorrelated log-normal relaxed clock and applied two calibration points. Firstly, we calibrated the crown of the Lamprologini to be 4 million years old (log-normal prior, mean of 4 Myr 95% conf interval: [3, 5]), based on the results from Meyer et al. (2016). Secondly, we included two riverine Lamprologini species *(L. congoensis* and *L. teugelsi*), and calibrated the onset of their branching event at 1.7 Ma (offset 1.1, log normal distribution with mean 1.7, 95% conf interval [1.15, 3.47], “use originate = true”), following Koblmuller (2010). We applied 1/X priors on the clock rates, and log-normal priors on the substitution rates. All other priors were left at their default setting. As tree model we used the birth death model. The used BEAST configuration file (the Beauti xml) can be found in the supplementary material.

We ran 10 independent STARBEAST MCMC chains, of 750M trees each. Each chain was verified to have ESS values of at least 100 for all parameters. The first 10M trees were pruned from these chains as burn-in and then they were combined (we used the species tree, rather than the individual gene trees) into one large chain (of 7400M trees). Chains were thinned by taking only each 5,000^th^ tree. Using TreeAnnotator (from the BEAST 2 suite) we constructed a Maximum Clade Credibility tree (using all 1.48M trees after thinning), storing the mean heights.

We then pruned the tree from riverine species to obtain the pure *Lamprologini* tree on which we fitted our model. Instead of performing one ABC-SMC inference on the obtained MCC tree using a huge number of particles, which would be more accurate but computationally extremely demanding, we performed 100 parallel inferences using 10,000 particles each. We report the mean Bayes factor across these replicates.

#### Branching time uncertainty in the empirical tree

To account for uncertainty in the estimates of branching times in the *Lamprologini* tree we sampled 100 trees from the posterior distribution obtained by *BEAST. Sampling was performed at random, irrespective of the likelihood of the trees. In the Supplementary material we show that the distribution of summary statistics of the subset of 100 trees is similar to the distribution of the thinned chain. The 100 sampled trees were, like the Maximum Clade Credibility tree, also pruned to remove the riverine taxa and stored separately. For all 100 trees we performed both the ABC-SMC model selection algorithm and the ABC-SMC parameter estimation algorithm, to determine the impact of different branching times on the inferred water level model and associated parameters, and to determine whether the MCC tree is a good representation of the underlying variability.

## RESULTS

### Lamprologini *phylogeny*

The onset of diversification within the *Lamprologini* is estimated to be around 3.96 Ma (95% Highest Posterior Density interval (HPD): [3.09, 4.91]), which is very close to the prior we put on the node age, based on previous estimates (Meyer et al. 2016). Furthermore, we estimate the branching off of the Congo species (*N. congoensis* and *N. teugelsi*) from the *Lamprologini* in Lake Tanganyika to have occurred around 1.35 Ma (HPD: [1.12, 1.68]), which is a bit younger than previously obtained estimates (1.70 Ma, (Sturmbauer et al. 2010)). The topology of the Maximum Clade Credibility tree is largely consistent with previous findings (Sturmbauer et al. 2010) (Figure 1). Placement of *Neolamprologus fasciatus* as a close relative to *N. wauthioni* seems to re-iterate previously published evidence for introgressive hybridization (Koblmüller et al. 2007). For the three species not previously included in the *Lamprologini* phylogeny, *Lepidiolamprologus mimicus* was placed as a close relative to the other species within the genus *Lepidiolamprologus*, *Chalinochromis cyanophleps* was placed as a sister species to *Chalinochromis brichardi*, within the group of *Chalinochromis* and *Julidochromis* species, and in agreement with previous analysis (Kullander et al. 2014b). In contrast to previous findings (Kullander et al. 2014b), *Neolamprologus timidus* is not placed as a sister species to *Neolamprologus furcifer*, but rather associates with *N. mondabu* and *N. falcicula.* Again, in contrast to other previous findings (Gante et al. 2016), we place *N. olivaceous* outside the *brichardi* complex, which includes the model system species *N. brichardi* and *N. pulcher*. (but see the DensiTree representation, which shows that this is not true for all trees). Interestingly, we also do not infer *N. savoryi* to be phylogenetically clustered within the *brichardi* complex (the ‘Princess cichlids’ (Gante et al. 2016)), in contrast to Gante *et al.* (2016). We should take into account however that the analysis by Gante *et al.* is based on on full genome sequences from only a small group of species, in contrast to the limited number of markers from a large number of species that we used.

As a reference we inferred speciation and extinction using the constant-rates birth-death model (Nee et al. 1994). Using Maximum Likelihood (the function bd_ML in the DDD package (Etienne et al. 2012)), we obtained an estimate of 1.871 myr^-1^ for the speciation rate, and an estimate of 0.993 myr^-1^ for the extinction rate, for the MCC tree. We find an estimate of 0.87 myr^-1^ for the diversification rate (speciation - extinction) and an estimate of 0.531 myr^-1^ for the turnover rate (extinction / speciation). For the 100 trees sampled from the full chain, we obtain estimates of 3.02 myr^-1^ (95% HPD: [1.608, 4.947]) for the speciation rate and 2.409 myr^-1^ (95% HPD: [0.884, 4.530]) for the extinction rate. This translates into estimates of 0.61 myr^-1^ (95% HPD: [0.313, 0.985]) for the diversification rate, and 0.765 myr^−1^ (95% HPD: [0.518, 0.930]) for the turnover rate. Estimates for the birth-death model obtained during reconstruction of the tree using BEAST indicate an estimate of 0.864 myr^-1^ (95% HPD: [0.287, 1.459]) for the speciation rate and an estimate of 0.613 myr^-1^ (95% HPD: [0.181, 0.953]) for the relative death rate, which translates into an estimate for the extinction rate of 0.52 myr^-1^ (95% HPD: [0.156, 0.823]) per million years. This yields estimates of 0.334 myr^-1^ and 0.613 myr^-1^for the diversification and turnover rate respectively. The BEAST inferences include the riverine species, so speciation and extinction rates are expected to be a bit different.

### Parameter estimation

We estimated parameter values for the three models for all 100 trees sampled from the posterior. We report the parameter values across the combined posterior across all 100 trees. Note that variation in the parameter estimates results from two sources of variation: branching time variation across the 100 trees, and secondly variation in the parameter estimate within each ABC-SMC inference.

The model without water level changes is identical to the constant-rates birth-death model, and we find that our ABC-SMC estimates for sympatric speciation at high water level (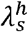) are slightly lower than the Maximum Likelihood estimate of the birth rate under the constant rates birth-death model (2.644 myr^-1^(95% HPD: [1.208, 4.633], see also Table 1) versus 3.02, see also Table 1). Similarly, we infer the extinction rate (μ) to also be slightly lower (1.950 myr^-1^ (95% HPD: [0.188, 4.101]) versus 2.409, see also Table 1). We obtain estimates of 0.694 and 0.738 for diversification and turnover respectively, which are close to the estimates obtained using Maximum Likelihood (a diversification rate of 0.610 and a turnover rate of 0.765 respectively). Taking into account the 95% confidence intervals on the obtained parameter estimates and the fact that the ABC-SMC estimates are potentially affected by the prior while the ML estimates are not, we are confident that estimates obtained using our ABC-SMC method for the model without water level changes are consistent with the maximum likelihood estimates under the constant-rates birth-death model.

**Table 1.**
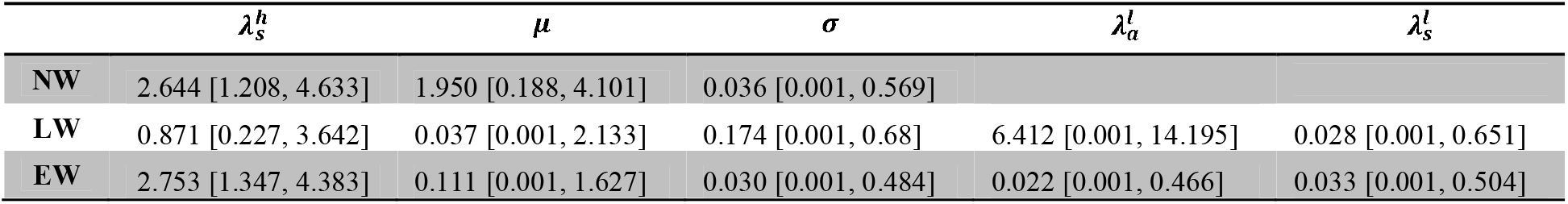
Median posterior density estimate, for sympatric speciation at high water 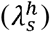, extinction (μ), perturbation (*σ*), sympatric speciation at low water 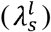 and allopatric speciation 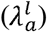. Shown are results for the model with no water level changes (NW), literature values for water level changes (LW) and water level changes extrapolated beyond the literature range (EW). The 95% credibility interval is shown between square brackets. All values are rates per million years.

Using the LW model, which implements water level changes following the literature (e.g. high water level until ~1.1 Ma, after which a series of water level changes took place), we infer a lower rate of sympatric speciation at high water level (0.871 myr^-1^ (95% HPD: [0.227, 3.642])), which is compensated with a high rate of allopatric speciation (6.412 myr^-1^ (95% HPD: [0.001, 14.195])) but not with a high rate of sympatric speciation at low water level (0.028 myr^-1^ (95% HPD: [0.001, 0.651])), suggesting that water level dynamics are important drivers of biodiversity, but only through allopatric speciation. Extinction is inferred to be low (0.037 myr^-1^ (95% HPD: [0.001, 2.133])). Because of the non-trivial relationship between speciation at high and low water level, we can no longer calculate diversification and turnover rates.

Using the EW model, where water level changes are extrapolated beyond 1.1 Ma, we observe that the rate of sympatric speciation at high water level is inferred to be similar to without water level changes (2.753 myr^-1^ (95% HPD: [1.347, 4.383])). Extinction, however, is lower (0.111 myr^-1^ (95% HPD: [0.001, 1.627])), and allopatric speciation and sympatric speciation at low water level are both inferred to be much lower than for the literature water scenario (0.022 myr^-1^ (95% HPD: [0.001, 0.466]) and 0.033 myr^-1^ (95% HPD: [0.001, 0.504]) respectively).

Across the three water level models we observe that the distribution of the post-hoc perturbations does not differ substantially from the prior for the NW and EW water models, with low estimates (0.036 (95% HPD: [0.001, 0.569]) and 0.030 (95% HPD: [0.001, 0.484]) for the NW and EW model respectively, Table 1). We notice a much higher value of associated with LW (0.174, (95% HPD: [0.001, 0.680])), which also has a much higher estimate for allopatric speciation at low water level. Allopatric speciation at low water level potentially causes temporal alignment of branching times and we introduced the parameter *σ* to correct simulated phylogenies for this, to allow comparison with phylogenies generated by *BEAST, which does not allow for temporally aligned branching times. Hence, the higher inferred value of *σ* for the LW model confirms the validity of the application of our post-hoc perturbation.

**Figure 3.**
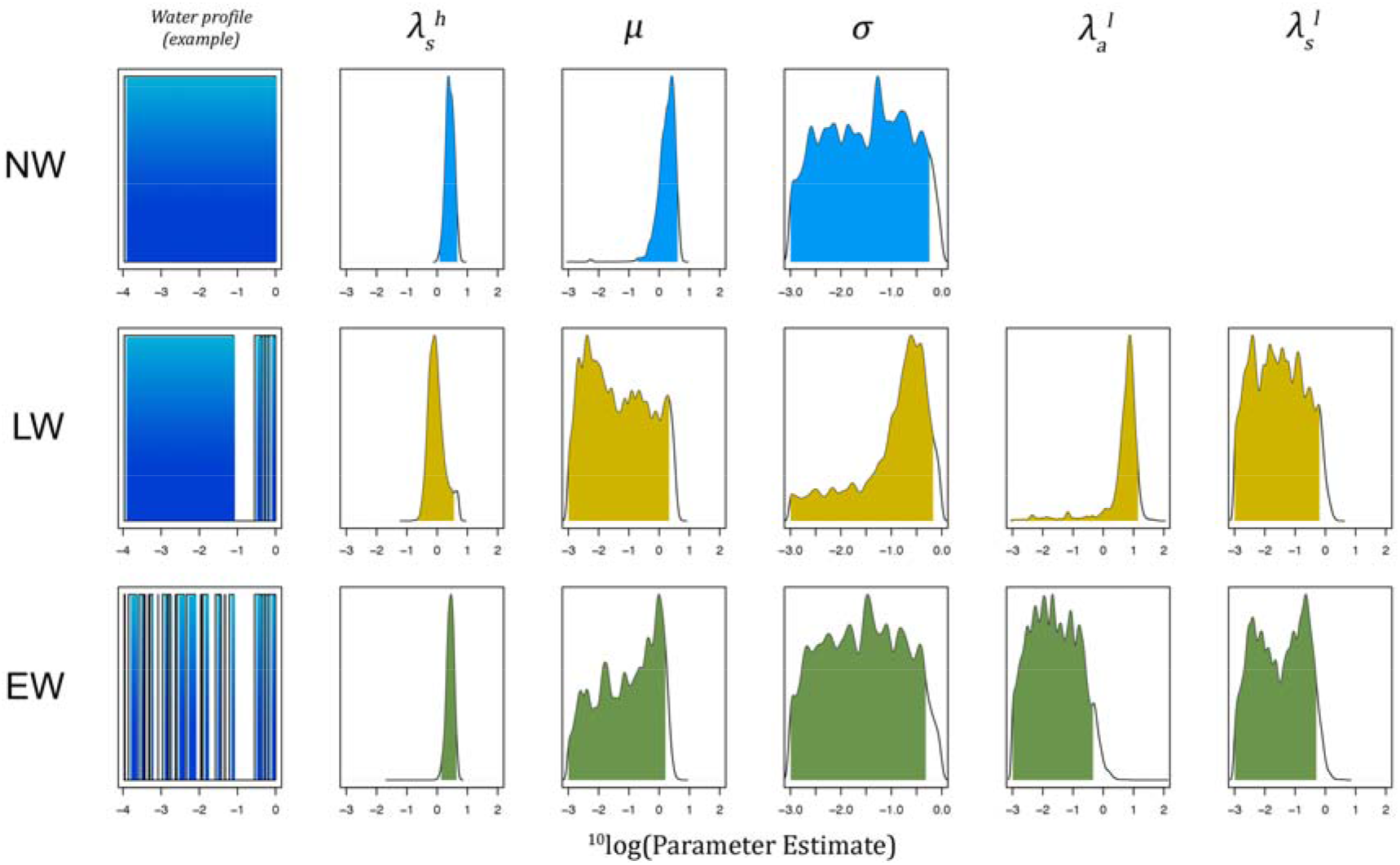
Posterior densities of the pooled posterior distribution across 100 randomly drawn trees from the posterior MCMC chain. Shown are estimates for the three water level scenarios (no water level changes (NW), literature values for water level changes (LW) and extrapolated values for water level changes (EW)). Shown are the posterior density (black line) and the 95% credibility interval (shaded area, blue for NW, gold for LW and green for EW. X-axes are on a ^10^log scale. The first column shows a sample water level profile, with the water level on the y-axis, and the time before present (in million years) on the x-axis. Note that for the EW model, for each simulation a new profile was generated, and that the shown profile is only one example of such a profile. Because without water level changes, 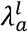 and 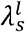 have no meaning, their posterior distribution is not shown for the NW scenario.

**Figure 4.**
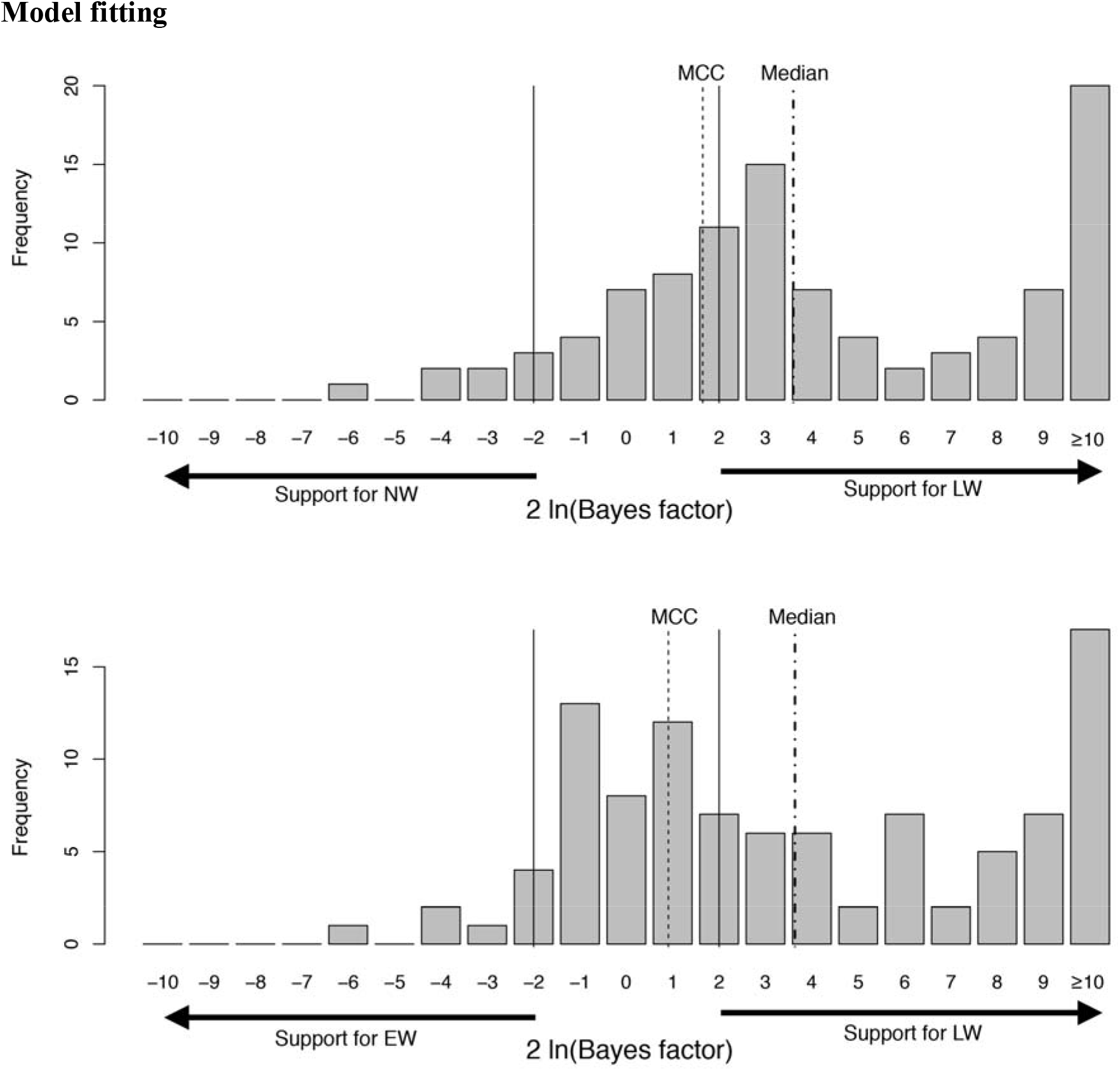
Model selection results on 100 trees randomly drawn from the *BEAST posterior of the *Lamprologini* tree. The top row shows 2 ln (Bayes factors) comparing posterior support of the LW (literature water changes) model with the NW (no water level changes) model, the bottom row shows 2 ln (Bayes factors) of the comparison between the posterior support for the LW model with the EW (extrapolated water level changes) model. A 2ln(Bayes factor) higher than 2 is generally considered to provide substantial evidence in favor of the respective model (Kass and Raftery 1995), which is indicated by the solid lines. The thin dotted line indicates the median 2 ln (Bayes factor) obtained for the MCC tree, for which we do not find substantial support for any of the three models. The thick dotted line indicates the median 2 ln (Bayes factor) for the trees drawn from the *BEAST posterior (e.g. the median of the distribution shown), which is in both cases above 2, indicating substantial support for the LW model compared to the other two models. 2 ln(Bayes factors) higher than 10 are grouped together into one category.

#### Model selection

When we apply the model selection algorithm to the MCC tree, we find median Bayes factors (we report here not the raw numbers, but 2 ln(Bayes factor), but for brevity refer to them as Bayes factors) of 1.64 and 0.9 when comparing the LW model with the NW and EW model respectively. We thus find no convincing evidence for any of the three models, when fitting our model to the MCC tree. Alternatively, when we fit to 100 trees randomly sampled from the *BEAST posterior, we find Bayes factors of 3.60 and 3.65 when comparing LW model with the NW and EW model respectively. Furthermore, in 77 out of the 100 trees we select the LW model as the most likely model (based on the Bayes factor), in 17 out of 100 trees we select the EW model, and only in 6 out of 100 trees we select the model without any water level changes.

**Figure 5.**
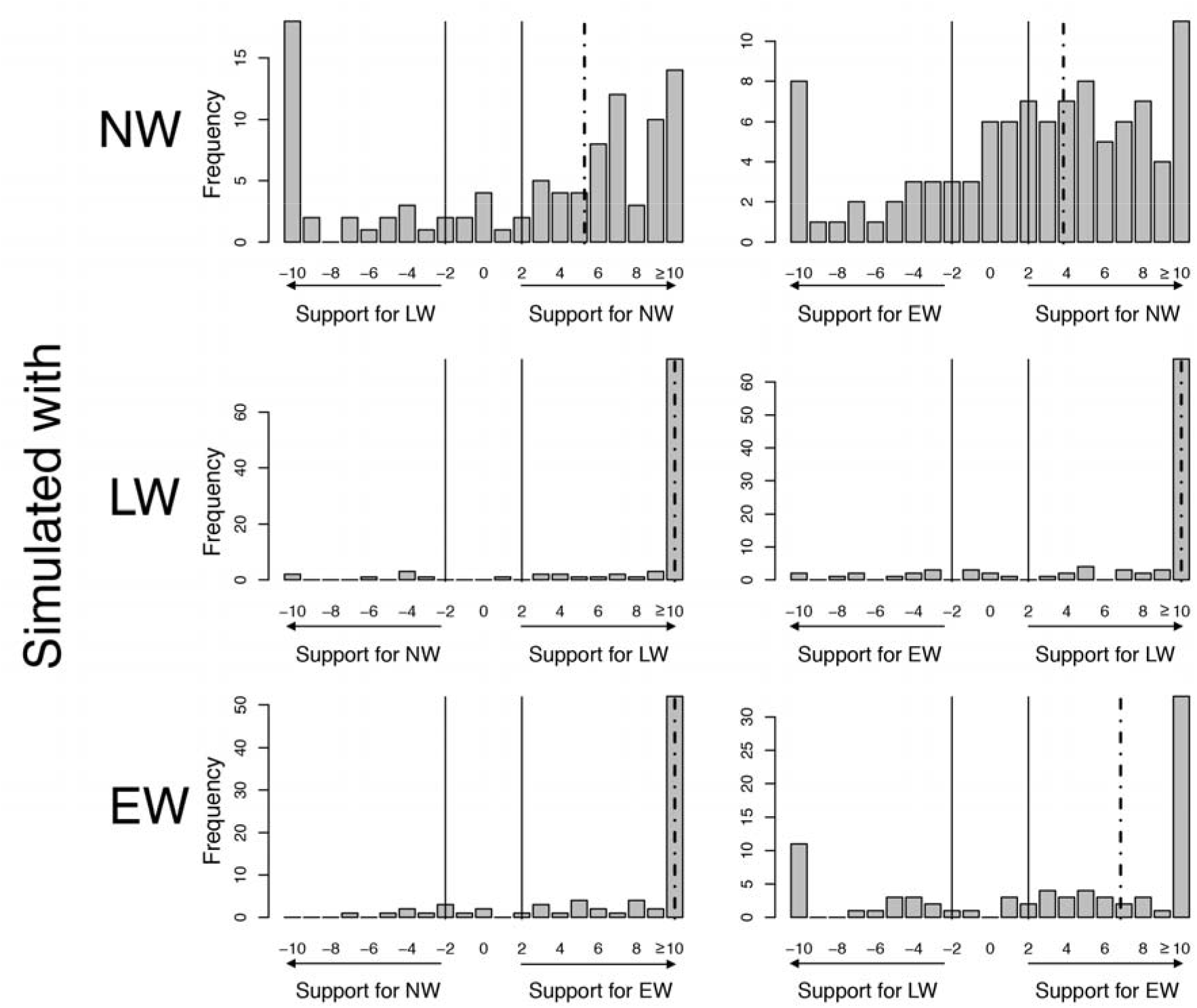
Validation of the ability of our ABC-SMC algorithm to infer the correct model. 100 replicate datasets were generated for each water level model (no water level changes NW, water level changes from the literature LW, or water level changes extrapolated beyond the literature range, EW). The plots show the distribution of the 2 ln(Bayes factor) across all 100 replicate inferences. The dotted line indicates the median 2 ln(Bayes factor). A 2ln(Bayes factor) higher than 2 is generally considered to provide substantial evidence in favor of the respective model (Kass and Raftery 1995). 2 ln(Bayes factors) higher than 10 are grouped together into one category.

Model validation shows that when we simulated data using the NW model, the NW model was selected using our model validation algorithm more than the other two models (59 out of 100 replicates). Median Bayes factors are both higher than 2, with a median of 5.28 and 3.83 versus the LW and EW model respectively, supporting considerable support for the NW model over the other two models. When data was simulated with the LW model, we selected the correct model in the majority of 100 replicates (84 out of 100 replicates). The Bayes factors reflect this, with medians of 18.4 (this is the maximum score) versus both the NW and EW model. Lastly, when we simulated data using the EW model, we selected the correct model more than the other two models, in 65 out of 100 replicates. This was reflected by the Bayes factors as well, as the median Bayes factor versus the NW model was 18.4, and the median Bayes factor versus the LW model was 7.47.

More interesting is the correct detection rate of a model, which is given by the number of trees simulated by the model that is selected for that tree. This is equal to asking whether, given posterior support for a respective model, we also find that the tree for which we find this support was simulated with the respective model. If our model selection procedure can not detect models accurately, we expect a detection rate of around 50%, as support is always divided between two (not three) models. Detection rates larger than 50% support the conclusion that our model selection procedure can adequately infer the correct model.

We find that across the 300 simulated trees, 120 trees received considerable support for the LW model over the NW model (e.g. 2 ln (BF LW/NW) > 2), of these 120 trees, 92 trees were simulated with the LW model, which leads to a correct detection rate of 77% (See Figure 6). Furthermore, out of 107 trees that received considerable support for the LW model over the EW model, we find that 83 trees were simulated using the LW model, which translates to a detection rate of 78%. We find similar detection rates for the NW model: 90% against the LW model (62 out of 69 detected trees) and 83% against the EW model (54 out of 65 detected trees). Lastly, detection rates for the EW model mirror these findings: a detection rate of 79% against the NW model (79 out of 100 detected trees), and of 86% against the LW model (68 out of 79 detected trees).

**Figure 6.**
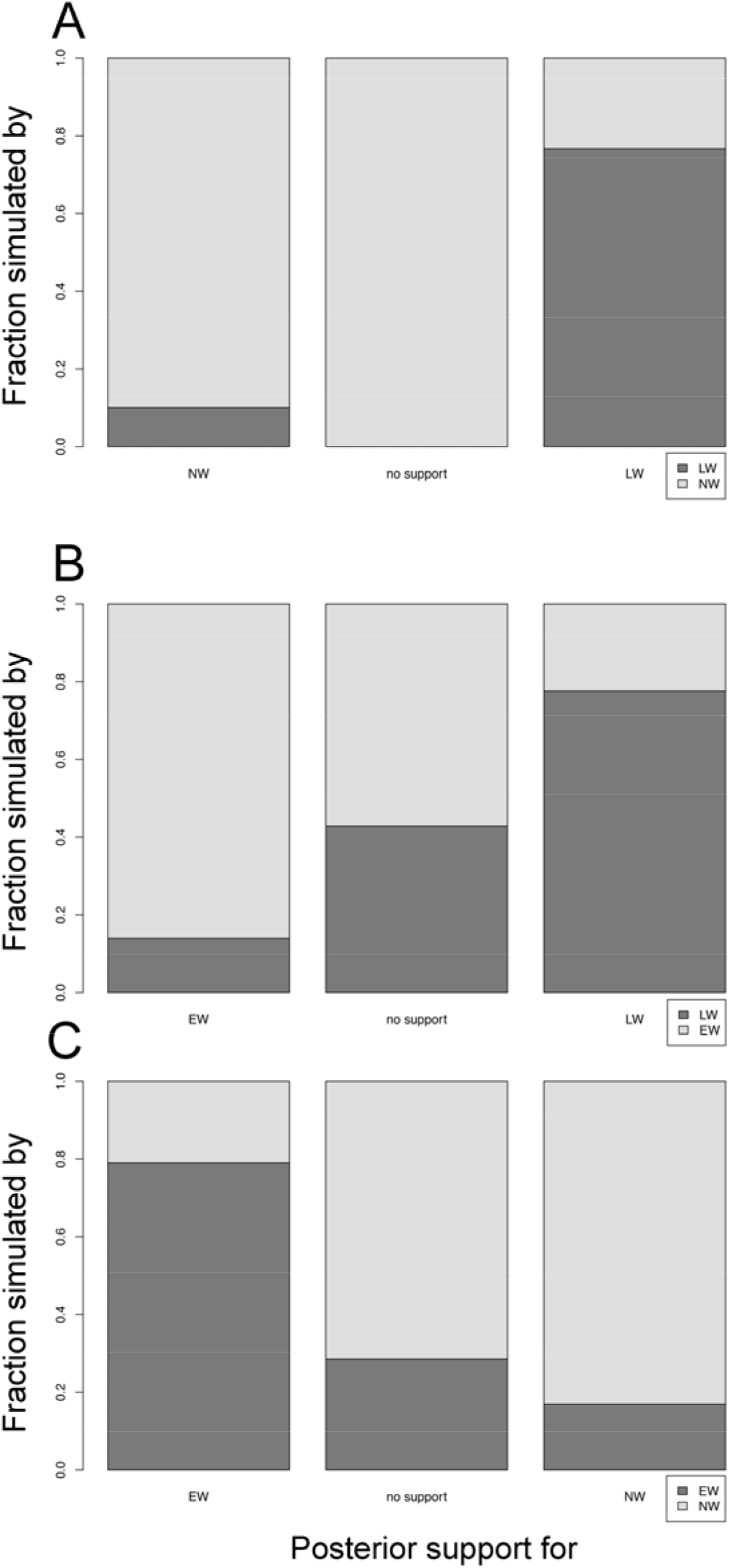
Accuracy of assignment of models depending on their posterior support. **A**: The relative fraction of trees simulated with either LW (dark) or NW (light), receiving support for LW (2ln(Bayes Factor LW/NW) > 2), support for NW (2ln(Bayes Factor LW/NW) < -2), or receiving no support for either model. **B**: The relative fraction of trees simulated with either LW (dark) or EW (light), receiving support for LW (2ln(Bayes Factor LW/EW) > 2), support for EW (2ln(Bayes Factor LW/EW) < -2), or receiving no support for either model. **C**: The relative fraction of trees simulated with either NW (dark) or EW (light), receiving support for NW (2ln(Bayes Factor NW/EW) > 2), support for EW (2ln(Bayes Factor NW/EW) < -2), or receiving no support for either model.

## DISCUSSION

We have presented a model that infers past speciation and extinction rates, and their interactions with changes in the environment, from a given phylogeny. We have shown that our model is able to accurately select between different scenarios, including or excluding environmental change. We applied our model to an updated phylogeny of the cichlid fish tribe of *Lamprologini* and found evidence that past water level changes have shaped current cichlid diversity in Lake Tanganyika, when we applied our model to a sample from the posterior distribution of trees of the *Lamprologini*, as inferred by *BEAST. We asked the model to select the best fitting of three scenarios: a scenario without any water level changes, a scenario using the values found in the literature, and a scenario using the mean rate of water level change found in the literature to extrapolate water level changes beyond the range of literature values available. We found that the model following literature water levels received most support, which suggests that water level changes have been an important driver of diversity in the *Lamprologini*. We note that a model without effect of water level changes on diversification (NW) can sometimes generate patterns that resemble the predictions of the preferred model (LW). Yet, we find when fitting our model to trees drawn from the *BEAST posterior that the distribution of Bayes Factors is skewed towards the model following literature water levels (LW) and we find support for the model without an effect of water level changes on diversification (NW) only for a small number of trees, suggesting that this effect is relatively small.

When we applied our model selection algorithm on the Most Credible Consensus (MCC) tree, we found contrasting results. Support for both models including water level changes diminished, and posterior support for the model without any water level changes increased. Nevertheless, no single model could yield enough support to convincingly reject the other two. Moreover, results using the MCC tree are markedly different from those using trees sampled from the posterior. We conclude therefore that the MCC tree, at least for the *Lamprologini*, but most likely more generally, provides a poor summary of the true species tree and of the underlying variation in branching patterns. Hence, we suggest to avoid reporting MCC trees, and instead to provide the reader with the full posterior distribution, for instance through a DensiTree plot (Bouckaert and Heled 2014). Posterior inference, for instance of speciation and extinction rates should preferentially also be performed on multiple independent samples from the posterior, rather than on the MCC tree, as the underlying variation might lead to very different results, as we have shown here.

Discrepancies between the MCC tree and the posterior distribution of trees could also potentially clarify previously recovered inconsistencies when studying diversification, for example in shrews in the Philippines. The Philippines have been subject to strong sea level fluctuations, causing the fission and fusion of several islands, primarily during the Pleistocene (Brown et al. 2013). Population genetic evidence has convincingly shown that the location of such fused islands correlates strongly with genetic divergence between populations in many different species (Evans et al. 2003; Linkem et al. 2010; Siler et al. 2010; Oaks et al. 2013). Phylogenetic analysis however, has failed to show any evidence of diversification associated with Pleistocene water level changes (Esselstyn and Brown 2009). The basis for this phylogenetic analysis however, was an MCC tree. Repeating the analysis on the posterior distribution underlying the MCC tree could mitigate these problems, and could clarify the impact of Pleistocene water level changes on diversification in the Philippines archipelago.

When allopatric speciation rates are high, the resulting phylogenetic trees have internal nodes that have synchronized branching times, e.g. branching times that align with episodes of water level change. Although Phylogenetic reconstruction software is able to infer simultaneous branching events, it typically uses only two parameters (birth and death) to infer all branching events of the tree. Therefore, if it can accommodate the simultaneous events, it is unlikely to fit well to the non-simultaneous events, and vice-versa. Our finding of evidence for a substantial role of habitat dynamics in diversification can therefore be regarded as conservative. To improve the fit of trees generated by our model with trees generated by *BEAST we included an *a posteriori* perturbation parameter in our model. This parameter determines the standard deviation of a Gaussian perturbation kernel that is applied to each node after the simulation has completed. By perturbing each node, we minimized the probability that branching times align in time. We found that standard deviation increased in size with an increase in allopatric speciation, as expected. A less *ad hoc* solution to deal with the alignment of branching times in the tree would be to incorporate the model presented here as a tree prior in phylogenetic reconstruction software. Although this need not introduce any significant differences in the tree topology, the distribution of branching times could be substantially influenced, and any subsequent inference focusing on such patterns could be very different. Including such models in tree reconstruction software may require incorporation of ABC methods, and will be extremely computationally demanding, but our results justify such an endeavor.

Given that water level changes are only prevalent during the last million years before present, we cannot exclude the possibility that increased diversification due to reasons other than changing water levels has driven diversification during this period. On average, the LW model could be represented by a simple birth-death model with a rate shift around one million years ago. We expect however that although such a model could accommodate the increased average diversification, it cannot replicate temporal alignment in branching events due to water level changes. To examine this in more detail, we fitted a simple birth-death model with a rate shift around one million years ago to the trees obtained from the posterior (see Supplementary Information). In the absence of a likelihood for the LW model, we compared the nLTT statistic for the rate-shift model with that of the LW model, as the nLTT statistic should be sensitive to detecting temporal alignment of branching events, We find that our model is much closer to the empirical data than the rate shift model. We attempted to improve the fit of the rate-shift model by allowing the speciation rate in the model to shift up and down in line with the literature values of the water level changes. The two rates inferred by the model then represent speciation at low, and at high water level respectively. Although we do find an increase in the rate of speciation at low water level, the fit of this rate-shift model is still worse than that of the LW model. This supports our conclusion that water level changes influence the phylogeny not only through an increased speciation rate, but also through temporal alignment of branching times.

Although we refer in our model to the different implementations of speciation as sympatric and allopatric, care should be taken in interpreting these forms of speciation. We consider here allopatric speciation only on a large scale, where populations become allopatric over stretches of hundreds of kilometers (Sturmbauer et al. 2001). Large-scale isolation might not be necessary for cichlids, as some species can already be limited in gene flow by a sand stretch of 50 meters separating populations (Rico and Turner 2002). Such micro-allopatric speciation events are not captured by the allopatric speciation rate in our model. Rather, these local scale events are captured in our model by sympatric speciation. Hence, sympatric speciation in our model covers all degrees of speciation ranging from full sympatry to allopatry, providing that geographical isolation is smaller than that imposed by a water level change. Allopatric speciation in our model then solely refers to speciation events caused by geographical isolation over a large distance, driven by changes in water level, and inducing simultaneous branching events.

In our model we have assumed that when the water level drops, species distribute themselves equally over the two pockets of water that survive the water level drop. A more realistic model would allow for a skew towards one of the pockets, either dependent on the respective sizes of the pockets, the distribution of the species over the lake at high water level, or both. We have here refrained from including a parameter that regulates the distribution of species over the two pockets in order to avoid over fitting. Another possible extension of our model would lie into extending the approach towards three or more pockets, possibly combined with a parameter governing the distribution of species across these three pockets during a water level drop. Bathymetric maps of Lake Tanganyika suggest that for some water level changes it might split into three lakes (Coulter 1991). How a split of a species into three populations, and associated allopatric divergence and speciation, affects phylogenetic structure and affects temporal alignment in branching times remains currently unexplored and would be an interesting avenue for future work. Our results are strongly in line with population genomic analyses in a number of cichlid species including *Eretmodus cyanostictus* (Verheyen et al. 1996), *Tropheus moorii* (Koblmüller et al. 2011; Nevado et al. 2013; Sefc et al. 2017), *Variabilichromis moorii* (Nevado et al. 2013), *Altolamprologus* (Koblmüller et al. 2016) and *Telmatochromis temporalis* (Winkelmann et al. 2016), and resonate with population genomic findings across the three African Rift Lakes (Sturmbauer et al. 2001). Furthermore, population genetic studies have shown that water level fluctuations in Lake Malawi have been associated with population expansion in cichlid species (Arnegard et al. 1999; Sturmbauer et al. 2001; Genner et al. 2010), suggesting a potential role for water level changes in Lake Malawi as well. Phylogenetic reconstruction for Malawi cichlid species is problematic however, partially due to the young age of the species. However, considering that the geological record of Lake Malawi spans a much larger part of the total lifespan of the lake (Delvaux 1995; Lyons et al. 2015; Ivory et al. 2016) and thus provides a much better record about water level fluctuations since the colonization of the lake by cichlids, we expect that modern genetic developments will soon allow for a thorough understanding of the impact of water level changes on cichlids in Lake Malawi as well.

### Conclusion

Our model integrates standard constant-rate birth-death mechanics with environmental change and with speciation induced by geographical isolation. We analyzed the phylogeny of the tribe of *Lamprologini* to see whether past water level changes in Lake Tanganyika have contributed to the current diversity of cichlid fish in Lake Tanganyika. We find an important role for environmental changes in driving diversity, and find evidence that past water level changes have shaped current standing diversity in the tribe of *Lamprologini*. However, we found that inference of past environmental changes from a single phylogeny, and more specifically, from the MCC tree, tends to lead to unreliable results. We therefore advocate caution when using the MCC tree as a basis for further analysis. Furthermore, we argue for the inclusion of more detailed branching models in phylogenetic reconstruction software, which allow for the inclusion of an interaction between the environment, and speciation rates.

## Acknowledgements

We thank Lucas Molleman for useful discussions. We thank the Netherlands Organisation for Scientific Research (NWO) for financial support through VIDI and VICI grants awarded to RSE. We thank the Donald Smits Center for information Technology of the University of Groningen for their support and providing access to the Millipede and Peregrine high-performance computing cluster. We thank the Max Planck Institute for Evolutionary Theory for their support and providing access to their high-performance computing cluster. We thank the Carl von Ossietzky Universität Oldenburg for their support and providing access to the CARL computing cluster.

